# First Report of Recurrent Genomic Selection with Real Data in Popcorn and Genetic Gain Increases

**DOI:** 10.1101/466003

**Authors:** Ismael Albino Schwantes, Antônio Teixeira do Amaral, Janeo Eustáquio de Almeida Filho, Marcelo Vivas, Pablo Diego Silva Cabral, Amanda Gonçalves Guimarães, Fernando Higino de Lima e Silva, Pedro Henrique Araújo Diniz Santos, Messias Gonzaga Pereira, Alexandre Pio Viana, Guilherme Ferreira Pena, Fernando Rafael Alves Ferreira

## Abstract

Recurrent Selection increases the frequencies of favorable alleles for economically important traits, which in the case of popcorn are popping expansion and grain yield. However, is time-consuming, since each selection cycle consists of three stages: progeny development and evaluation, and recombination of the best families. With the Recurrent Genomic Selection use, the time required for each selection cycle can be shortened, as it allows the evaluation and recombination phases to be performed simultaneously, reducing the time needed to complete one selection cycle to only one growing season. In this respect, the objective of this study was to determine the selection accuracy and genetic gains for different selection strategies: PhEN = estimates based exclusively on the phenotypic data of 98 plants; PhEN + GEN = estimates based exclusively on the phenotypic and genotypic data of 98 plants; and GEN = estimates based exclusively on SNP marker genotyping. The following traits were evaluated: 100-grain weight, ear height, grain yield, popping expansion, plant height, and popcorn volume. Field trials were carried out with 98 S_1_ progenies, at two locations, in an incomplete block design with three replications. The parents of these progenies were genotyped with a panel of ~ 21K SNPs. From the results based on the predictions by strategy GEN, at different selection intensities, the average annual genetic gain for the different traits was 29.1% and 25.2% higher than that by the strategies PhEN and GEN + PhEN for 98 selection candidates; 148.3% and 140.9% higher for 500; and 187.9% and 179.4% higher for 1,000 selection candidates, respectively. Therefore, recurrent genomic selection may result in a high genetic gain, provided that: i) phenotyping is accurate; ii) selection intensity is explored by genotyping several plants, increasing the number of selection candidates, and iii) genomic selection is used for early selection in recurrent selection.

## Introduction

Recurrent selection (RS) is one of the most practical breeding methods in the development of popcorn maize varieties. The technique is used to increase the frequencies of favorable alleles for commercially interesting traits (1,2), namely popping expansion and grain yield, in the case of popcorn. In addition, it preserves the genetic variability, enabling new breeding cycles. However, it is a laborious and time-consuming procedure, since each selection cycle involves three stages: progeny development and then evaluation, and recombination of the best families. The evaluation and recombination stages are not carried out at the individual level, and therefore, families from progenies considered superior are used. This leads to a reduction in the selection gain per cycle. In addition, the period required to complete a RS cycle comprises three growing seasons, thus reducing the annual genetic gains (2).

By Recurrent Genomic Selection (RGS) - a procedure that applies the principles of Genomic Selection (GS) in recurrent selection - the time needed for each selection cycle can be shortened, since by RGS, the evaluation and recombination steps can be carried out simultaneously. This method is a form of direct early selection, i.e., it has a premature effect on genes expressed in adulthood (3,4). Consequently, the time required for one selection cycle is shortened to only one growing season.

In the literature, no studies on GS in real popcorn breeding populations have assessed the genetic gains for agronomic traits and the viability of this technique in comparison with traditional breeding techniques. However, GS has been proved to be efficient to increase selection gains per time unit and as a tool to predict effects of general combining ability of testcrosses and genotypic values of double-haploid maize lines (5); in the breeding of allogamous lines (6) and in the prediction of genotypic values of single-cross maize, wheat and rice hybrids (7,8).

In some cases, GS also proved to be more efficient per unit of time than phenotypic or marker-assisted selection (9). In addition, other studies also reported that GS is efficient and can be used in plant breeding (10–12).

Therefore, the objective of this study was to determine the selection accuracy for different selection strategies and the respective genetic gains at different selection intensities, with a view to assessing the efficiency of recurrent genomic selection in popcorn breeding.

## Material and Methods

### Study population, genotyping and phenotyping

The study was carried out with the popcorn population UENF-14, in the eighth cycle of Recurrent Selection (RS). After recombining the selected progenies, the seeds of the resulting population were sown, initiating the ninth RS cycle, which was used in this study. Two hundred plants were sampled for genotyping, of which 98 plants were selfed for the phenotypic evaluation of their S_1_ progenies.

Based on the 200 DNA samples collected from seedlings, the polymorphism of the genome among plants of this population was described by the Capture Seq method (13), using 5,000 well-distributed probes in the maize reference genome. This genomic approach was conducted in collaboration with the company Rapid Genomics LLC, resulting in 21,442 SNPs. Based on genotypic information, using software Plink (14), the three following sequential filters were applied: a) elimination of individuals with > 10% missing data b) SNP elimination of SNPs with > 5% missing; and c) elimination of SNPs with Minor Allele Frequency (MAF) <5%, resulting in 196 plants with 10,507 SNPs. Of the filtered individuals, 98 were used in the S_1_ progeny trial.

With these 196 plants and 10,507 SNPs, the population genome was characterized, estimating the linkage disequilibrium (LD) using r^2^ statistics with software Plink (14). Based on the LD results, an LD decay curve was fitted according to (15), using the nlm function of the R language. In addition, the kinship coefficient of the plants was calculated, obtaining the Genetic Relationship Matrix (GRM) with the R package rrBLUP (16), but adding a small value (10^−2^) diagonally to improve the numerical stability (17,18).

With the resulting S_1_ families, two trials were set up, one at the Colégio Estadual Agrícola Antônio Sarlo, in Campos dos Goytacazes, in the northern region of the State of Rio de Janeiro, (lat. 21° 45 S, long. 41° 20′ W; 11 m asl), classified as a tropical rainforest climate, with annual averages of 23° C, 1,023 mm rainfall, and potential annual evapotranspiration of 1,601 mm. The other experiment was installed at the Experimental Station PESAGRO-RIO, in Itaocara (lat. 21° 39’ 12’’ S, long. 42° 04’ 36’’ W; 60 m asl), with annual averages of 22.5° C and 1,041 mm rainfall. Both experiments were initiated in August 2016.

The experiment was arranged in an incomplete block design, with three replications. To this end, seeds of each family were sown in 5.0 m long rows, with a spacing of 0.9 m between rows and 0.2 m between plants, with three seeds per planting spot, at a depth of 0.05 m. At 21 days after emergence, thinning to one plant per spot was performed, resulting in a total density of 60,000 plants per hectare. Fertilization at planting was applied based on the soil chemical analysis and sidedressing about 30 days after planting. Management practices were used according to the crop demands.

The following phenotypic traits were measured: i) mean plant height (PH), i.e., the distance, in cm, from the soil level to the insertion of the flag leaf, after tasseling of six healthy plants; (ii) mean ear height (EH), i.e., the distance, in cm, from the soil level to the insertion of the first ear in six plants per plot; iii) 100-grain weight in grams (100GW); (iv) grain yield (GY), expressed in kg.ha^-1^; v) popping expansion (PE), determined in plastic containers without oil, with three replications per plot, using grain aliquots weighing 30 grams each. Popping was carried out in the microwave oven for 2 min, the expanded popcorn volume measured in a 2,000 mL beaker, the resulting value divided by the initial grain weight of 30 g and the final result expressed by the ratio mL g^-1^; and vi) expanded popcorn volume per hectare (PV), obtained by multiplying the mean yield per plot by popping expansion, generating the mean expanded popcorn volume per hectare of plantation, expressed in m^3^.ha^-1^ (PV = GY x PE/1000).

### Statistical method applied to genomic selection

This study used a model of individuals, in which the pedigree - based kinship matrix was replaced by a kinship matrix estimated by markers, whose model is called Genomic Best Linear Unbiased Predictor (GBLUP). The advantage of GBLUP is that the direct prediction of the Genomic Estimated Breeding Value (GEBV) of individuals is computationally much simpler, since the number of markers is usually far higher than that of individuals (19).

The GBLUP models are based on the Genetic Relationship Matrix-GRM matrix, obtained by marker panel data (20). This matrix was based on the assumption that every marker with the same MAF also contributes equally to the genetic variance. Thus, this model is consistent with that of negligible inheritance, traditionally assumed in classical quantitative genetics, in which the traits are assumedly polygenic, with a number of genes that tend to infinity, and all contribute equally to the phenotypic variation. According to the experimental conditions of this study, the GBLUP model was fitted as follows:

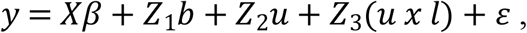

where: **y** is the vector of phenotypic data of a given trait; ***β*** the vector of fixed effects that include the model intercept, effect of replication within location, effect of location and co-variables as number of plants per plot, counted immediately after thinning and grain moisture, only for the traits 100GW, GY, PE and PV; ***b*** is the random effect of incomplete block within the replication and location; **u** is the random effect of the genomic additive genetic value (GEBV); (***u x l***) is the random effect of the interaction between GEBV and location; and is the random error effect.

For this model it was assumed that:

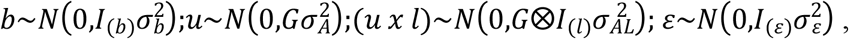

where ***I***_(***b***)_, ***I***_(***l***)_ and ***I***_(***ε***)_ are identity matrices with order equal to the length of vector **b**, number of locations and number of observations, respectively; **G** is the GRM matrix described above; ⊗ the Kronecker product; 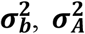 and 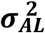 are the components of variance associated with the blocks, additive variance and variance component of the additive genetic interaction with the location, respectively. The variance components were estimated by REML with algorithm AI, using the statistical package ASReml (21) of software R (22).

The genetic and environmental correlation components were also estimated, based on bi-trait and bi-environment models,

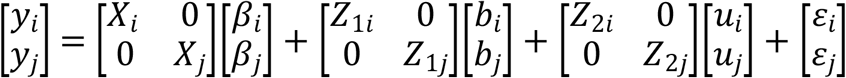

where the subscripts **i** and **j** represent two traces in the same environment, or the same trait in both environments, considering in this model that:

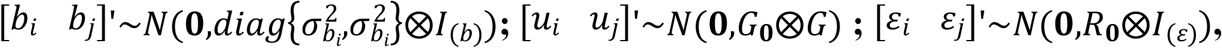

where the matrices 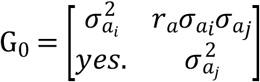 and 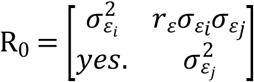. The other terms of the model are analogous to the uni-trait model.

### Estimates of selection accuracy and genetic gain

Selection using SNP genotyping will only be efficient if there is a considerable increase in the response to selection per time unit when compared to the selection based on family means, since the comparison of these means underlies the selection of the best progenies throughout the cycle of the recurrent selection program for UENF-14.

Thus, to estimate selection accuracy and gains, three different strategies were used, namely: i) PhEN = estimates based exclusively on the phenotypic data of 98 individuals; ii) PhEN + GEN = estimates derived from phenotypic and genotypic data of 98 individuals; and iii) GEN = estimates based exclusively on genotyping by SNP markers. The GBLUP model was adjusted with 90% of the individuals of the population, whereas the genetic merit of the remaining part (10%) was predicted based only on marker data. This model fitting process was repeated 10 times, and in each cycle, the genetic merit of a different group of individuals was predicted with the ignored phenotype. This process is known as ten-fold cross-validation (23,24), and was used in this study to test the efficacy of early selection, using a previously fitted model.

By the strategy GEN, predictions for different selection intensities were tested, for 98 to 1,000 selection candidates, setting an absolute value of 40 selected plants. Thereafter, a comparative evaluation of the annual genetic gain by the strategies GEN, PhEN and GEN + PhEN was carried out. Accuracy was estimated as proposed by (25) considering only the estimates of the 98 individuals that passed the filter for the loss rate of SNP data in this study.

The following formula was used to compute the genetic gain:

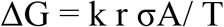

where: k = standardized selection differential, where a number of 40 selected plants was fixed for this study; r = selection accuracy, determined by the predictive accuracy estimated by each method; σA = square root of the additive variance, estimated based on the genotypic and phenotypic data; and T = time to complete a cycle; and for the strategies PhEN and GEN + PhEN, the progenies must be phenotyped and the normal duration of Recurrent Selection is three growing seasons (T = 1.5 years under tropical conditions), while in the case of strategy GEN, in which the phenotyping step is ignored, selection occurs at the plant level rather than that of progenies. As selection occurs before flowering, the selected plants are directly recombined so that one RS cycle with a previously fitted model requires a duration of one growing season (T = 0.5 year under tropical conditions). A period of 0.5 years for strategy GEN and of 1.5 years for GEN + PhEN was used, considering the possibility of consecutive growing seasons of recurrent selection. In many scenarios, the environment requires a 2-year period to complete the full cycle, due to the impossibility of cropping in the cold season.

## Results

### Study population and genetic parameters

To improve the understanding of some genetic traits of the study population, several parameters were estimated with the panel of available SNPs. After the filters for data loss, slightly more than 10% of SNPs had fixed alleles in the population, whereas ~ 25% proved to be polymorphic, but with low MAF (<0.05). Therefore, a satisfactory amount of ~ 65% of well-distributed genome markers have segregating alleles with high frequency in the study population (Figure 1a). The r^2^ between adjacent SNPs was in the mean ~ 0.36 (SD = 0.36), and half of the decay was estimated to be ~ 107.54 Kb (Figure 1b). The inbreeding coefficient based on markers is distributed around zero (Figure 1c), consistent with a population derived from a random recombination of plants, although the inbreeding coefficient of ~ 89% of these plants, resulting from eight previous RS cycles, was estimated between 0.1 and 0.3 (Figure 1d).

**Fig. 1.**
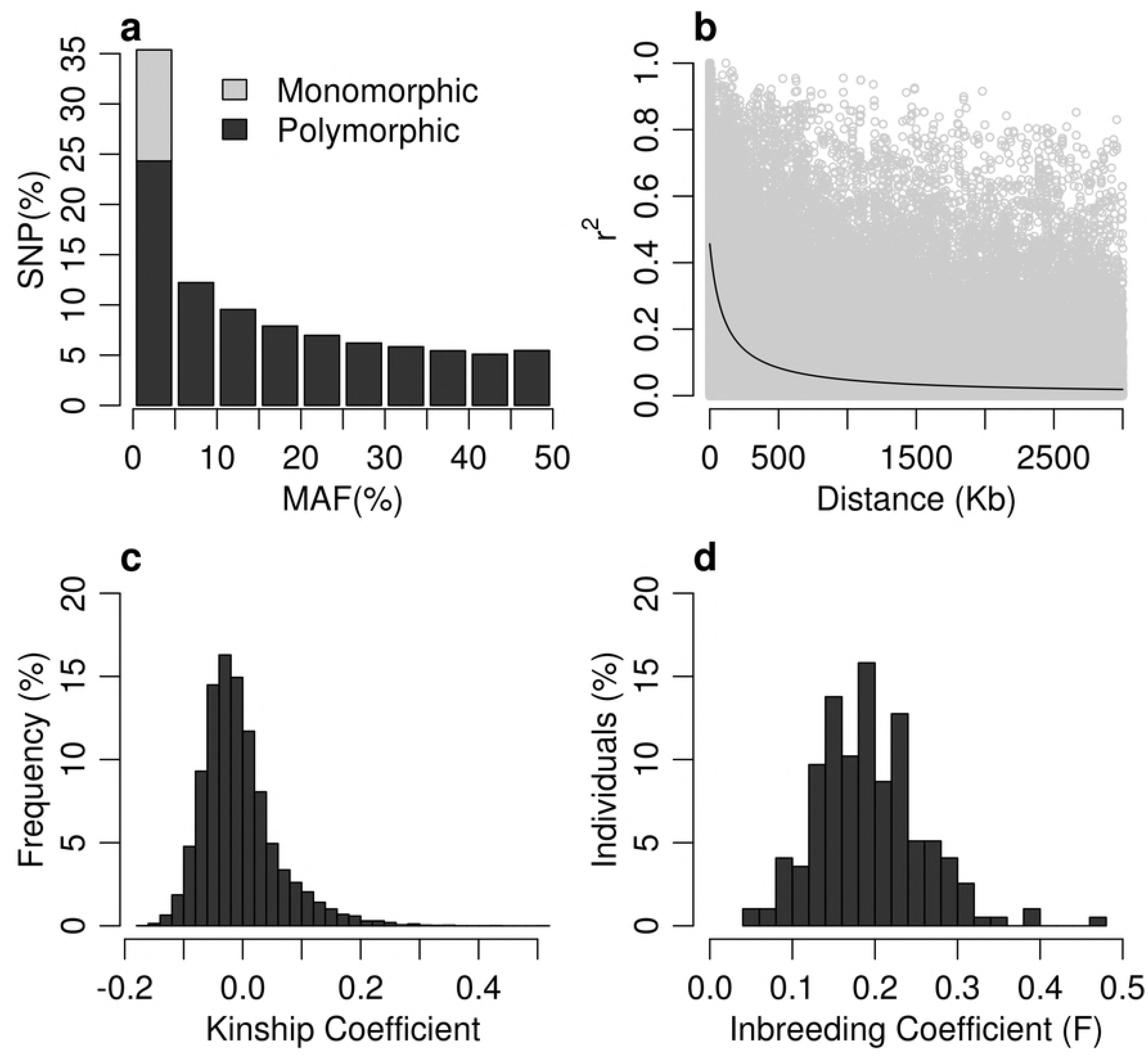
Genetic aspects of the study population. a) Distribution of allele frequencies; b) Linkage disequilibrium; c) Distribution of the kinship coefficient among plants; d) Distribution of the inbreeding coefficient of plants. Figure **a** was generated with a marker panel of 16,258 SNPs with < 5% missing data and Figures **b, c** and **d** were created with 10,507 SNPs of Figure a with MAF> 5%. In all figures, 196 plants with a data loss rate < 10% were considered.

The genetic variance component was significant for all evaluated traits at p <0.01. The variance of the genotype - location interaction was only significant for GY (p <0.01) and PV (p <0.01). The values of the coefficients of variation were acceptable for the studied traits (4).

Inheritability is one of the most relevant genetic parameters for breeding, due to its predictive capacity, expressing the confidence of the phenotypic value as a guide to the genetic value (2). It is therefore a key parameter in the choice of breeding strategies, since it influences decision making between selection techniques directly, making the development process of new cultivars more efficient (1).

In this study, the mean-based heritability was medium to high for most traits, and was < 0.5 only for PV and > 0.8 for 100GW, EH and PH (Table 1). For the main agronomic traits PE and GY, heritability was found to be 0.62 and 0.51, respectively. The correlation between heritability and accuracy was high for the different strategies (GEN = 0.99, PhEN + GEN = 0.93 and PhEN = 0.94), where higher heritability were associated with higher accuracy values.

**Table 1.**
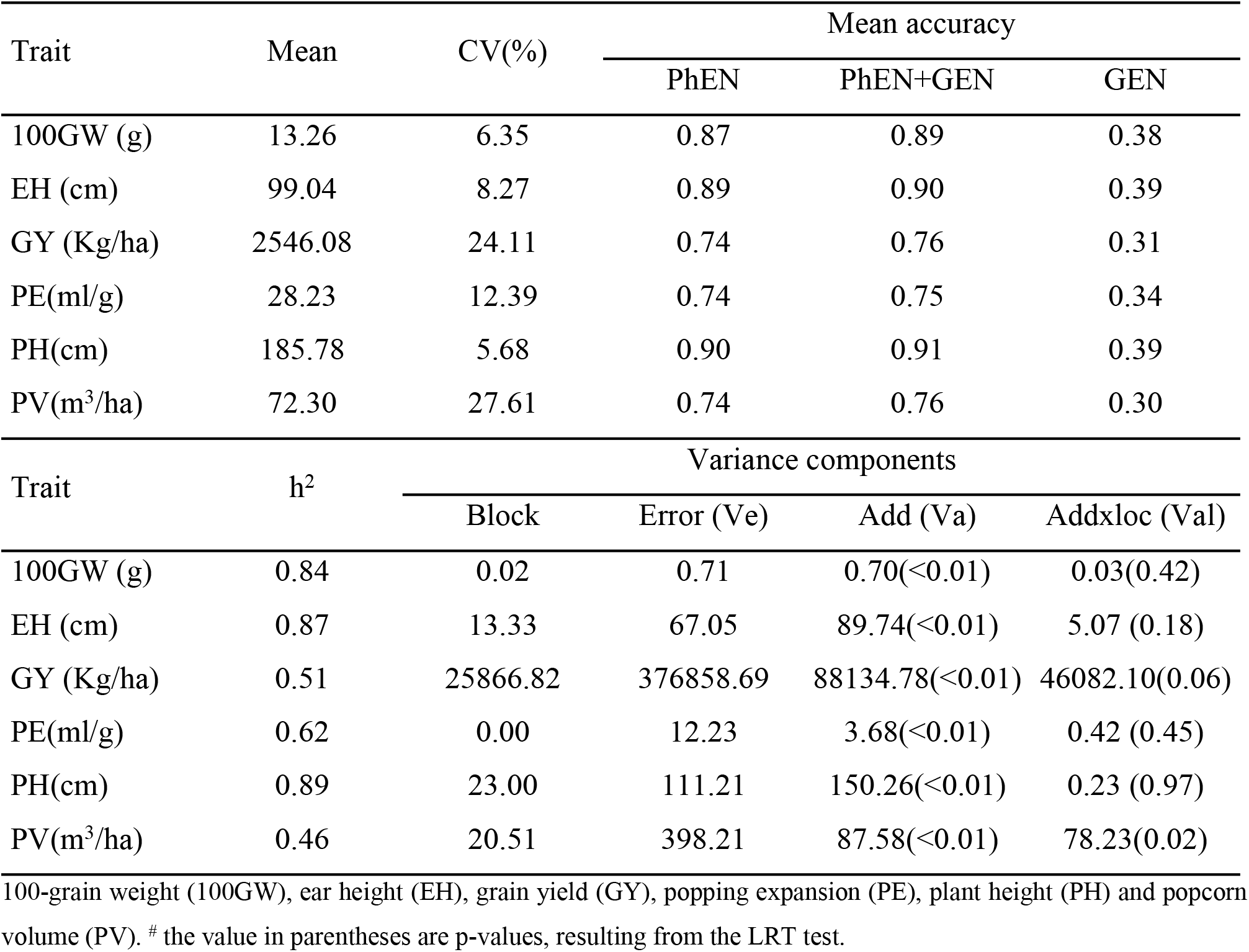
Selection accuracy estimates based on exclusively phenotypic data (PhEN), on phenotypic and marker data (PhEN+GEN) simultaneously and on exclusively marker data (GEN); components of additive variance (Va), interaction of additive-location effects (Val), error (Ve) and mean heritability (h²) for the evaluated traits.

### Accuracy of genomic selection

The accuracy of genomic selection is determined by the correlation between the estimated genomic genetic value and the genetic values resulting from phenotyping (26). Cross-validation is generally used to evaluate the accuracy of genomic selection (27). These validations are used to predict candidate phenotypes for future selections (28–30).

In conventional recurrent selection, selection is based on exclusively phenotypic data; in this study, when including information from the set of 10,507 SNPs markers, accuracy increased slightly for most traits. In this way, the model GEN + PhEN was in the mean ≈2.5% more accurate than the traditional model, which ignores the markers (PhEN). The accuracy based exclusively on the strategy GEN was between 0.25 and 0.39, i.e., much lower than the values based on PhEN and GEN + PhEN (Table 1).

The accuracy values for the different strategies were highest for the traits related to crop development, e.g., PH and EH. For the agronomically most important traits of popcorn (GY, PE and PV), accuracy values of > 0.74 were obtained by the strategies GEN + PhEN and PhEN. The accuracy and heritability values were also high for the secondary agronomic trait 100GW.

### Genetic and environmental correlations and genetic gain

The importance of simple correlation coefficients was highlighted by (1), since with these, the degree of genetic and non-genetic association between two or more traits can be quantified. The genetic correlation may be due to pleiotropy or gene linkage (in a population with linkage disequilibrium) among genes that are associated with two traits. Some genes may increase the phenotypic value of two traits, causing a positive correlation, while other genes can increase one and reduce the other, resulting in a negative correlation (31).

In this study, the genetic correlations between environments were high for most traits, with values close to 1, except for trait PV with a correlation of approximately 0.5. When considering both locations, the residual correlation was practically null for all traits (Figure 2).

**Fig. 2.**
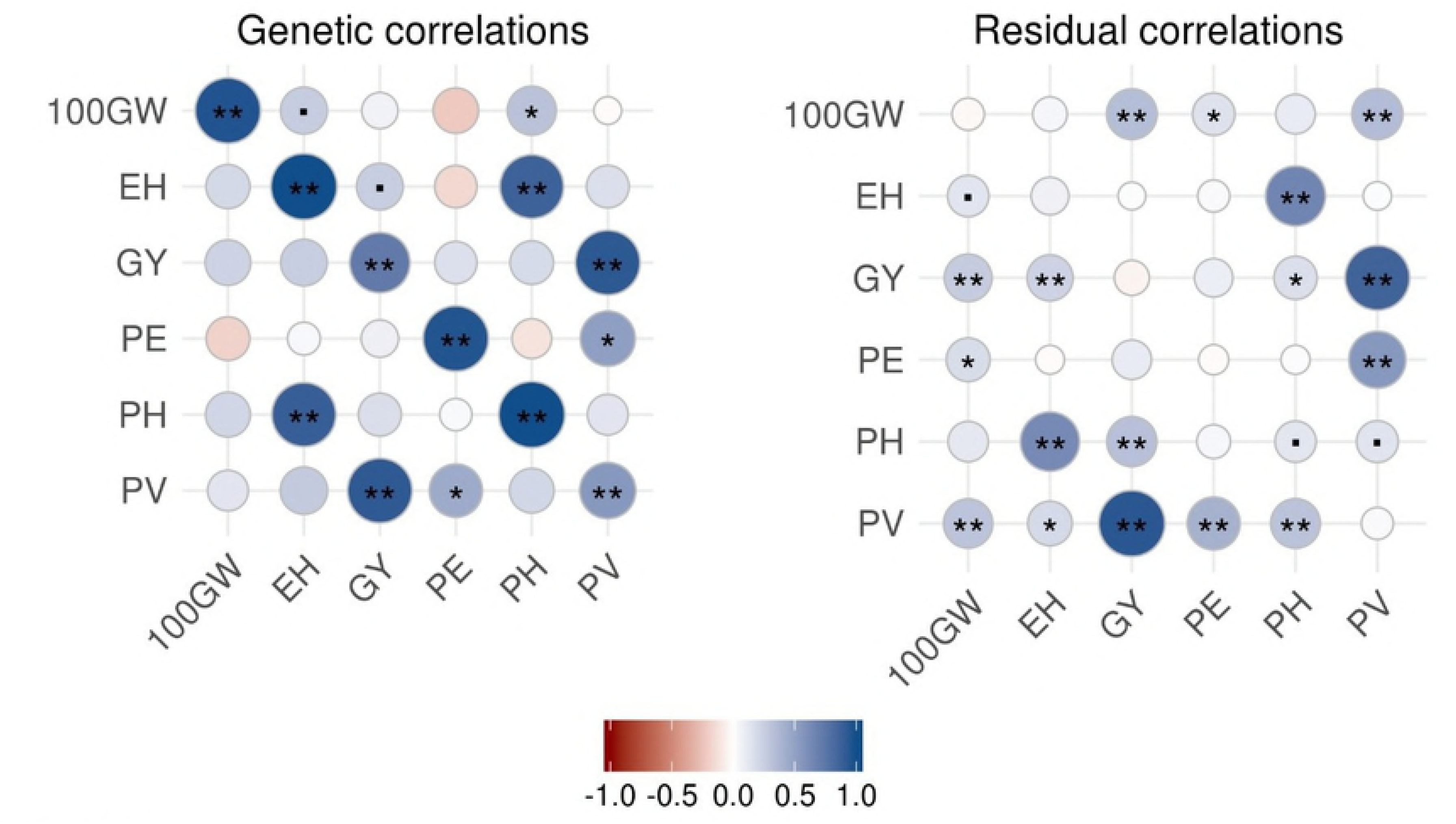
Diagrams of the genetic and residual correlations calculated for the traits 100-grain weight (100GW), ear height (EH), grain yield (GY), popping expansion (PE), plant height (PH), and popcorn volume (PV). Correlations between the traits, estimated in Campos dos Goytacazes and Itaocara, are located above and below the diagonal, respectively; the correlations of each trait between the locations are on the diagonals. **, * and dot “.”: stand for statistically different correlations from zero, at 1, 5 and 10% probability.

Among the observed genetic correlations, the trait pairs PH and EH; and GY and PV are highlighted, which were highly and positively correlated in both locations (Figure 2).

Genetic gain is the amount of increase in performance that is achieved in breeding programs over consecutive selection cycles (30). The selection gain depends on the selection differential, which in turn is the difference between the mean of the selected group and the mean of the original population. Therefore, in the selection process, the higher the selection pressure, the greater the differential, and consequently, the higher the genetic gain (2).

The results of predictions at different selection intensities by the strategy GEN showed that the average annual genetic gain for the different traits was 29.1% and 25.2% higher than by strategies PhEN and GEN + PhEN for 98 selection candidates; 148.3% and 140.9% higher for 500; and 187.9% and 179.4% higher for 1,000 selection candidates, respectively. Moreover, the average annual gain by strategy PhEN + GEN was 3% higher than by PhEN (Figure 3).

The strategy GEN obtained significant annual gains for the most important agronomic traits GY and PE, which were 32.8% and 28.2% higher for 98 candidates, and 155.4% and 147.1% higher for 500 candidates, respectively, than by strategy PhEN. By strategy GEN, the average annual gain increased 92.4% up to 500 selection candidates, and thereafter, the selection gain kept on increasing, although to a lesser extent, since for 500 to 1,000 selection candidates, the gain was 15.9% (Figure 3).

**Fig. 3.**
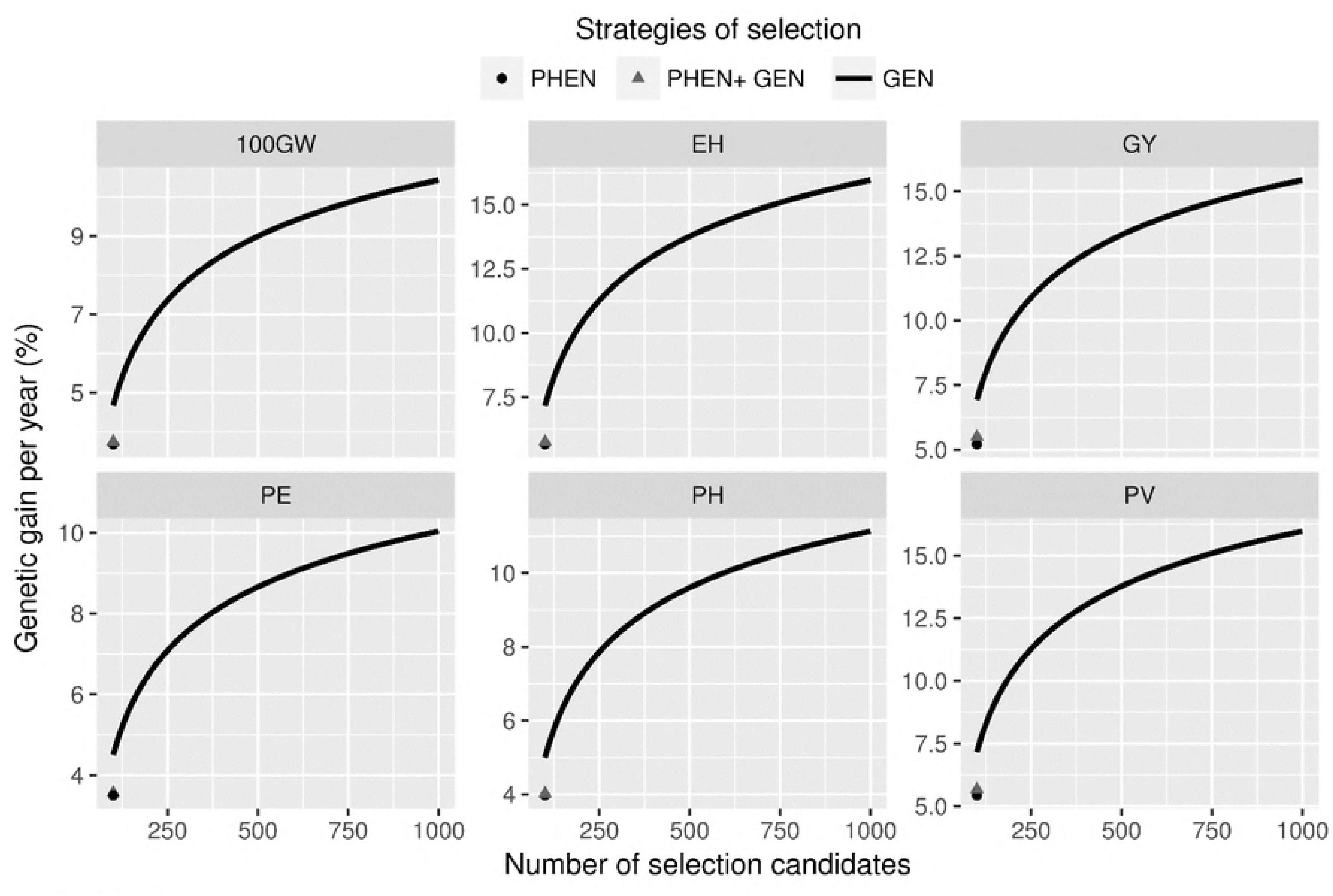
Annual genetic gain by the different tested strategies for the traits 100-grain weight (100GW), ear height (EH), grain yield (GY), popping expansion (PE), plant height (PH), and popcorn volume (PV). The tested selection strategies were based exclusively on data of the phenotypes (PhEN), on both phenotypic and SNP markers (PhEN + GEN) and finally, based exclusively on data of SNP markers.

## Discussion

### Genetic parameters

The significant genetic differences observed indicate the existence of genetic variability, enabling selection and genetic gains among the genotypes under study. Long-term genetic gains depend essentially on the potential genetic variability, which is maintained throughout the selection cycles and released by recombination at the end of each cycle. Thus, for the establishment of breeding populations, the genetic variability should be analyzed (32,33).

The variance of the significant genotype-location interaction found for the traits GY and PV is a result of differentiated responses of the tested genotypes in different environments. However, to infer whether the interaction is significant is only a qualitative approach, and the correlated response can only be investigated by the genetic correlation between the environments (34). For the trait GY, with a high genetic correlation, the selection in one environment will imply in a positive response in another. As the genetic correlation between the environments was positive, the selection based on the general additive value, i.e., for both environments, will provide a positive selection response for both environments.

The heritability of PE (0.62) exceeded that of GY (0.51). Also (35) observed a lower environmental influence on the expression of PE than on GY. The higher heritability of PE can also be explained by the fact that the trait is predominantly controlled by additive genes (36–39).

The low heritability (0.43) obtained with PV can be explained by the fact that this trait is a combination of GY and PE, also incorporating the environmental influence in the trait expression of two polygenic traits. According to (40), genomic selection will provide relevant genetic gains for plant breeding, even for low heritability traits, by reducing the duration of the selection cycle.

The high heritabilities (> 0.8) computed for the traits 100GW, EH and PH indicate the confidence of the transmission of the traits to future generations, directly influencing the selection gains (1). The high correlation of heritability with accuracy found agrees with results of (41–43). According to (43), when breeders outline the phenotyping experiments for GS, they should consider that the heritability of the traits of interest in the training population should be high to ensure a satisfactory predictive accuracy. To this end, the number of locations and replications should be increased.

### Prediction accuracy in genomic selection

By the strategy GEN + PhEN, the accuracy was higher for most evaluated traits than by PhEN, due to the increase caused by the use of the real kinship matrix, in other words, based on the markers of the individuals and not, as by the traditional approach, on a mathematical expectation (44).

The accuracy values by the strategy GEN were substantially lower than by the models PhEN and GEN + PhEN for all traits. Likewise, other studies reported that genetic gain prediction, considering only kinship information - based on genealogy and/or markers - showing that accuracy is lower in the absence of phenotypic data (45–47). Thus, to apply GS in breeding, other components of genetic gain should naturally be explored, e.g., selection intensity and a reduction in the intergenerational interval, since accurate phenotyping alone ensures reasonable selection accuracy.

The higher accuracy values for the crop development trait PH than for the agronomically relevant trait GY by the different strategies agree with results of (43) in maize. For trait PE, (12) identified an accuracy of 0.55 for a heritability of 0.3; and accuracy of 0.84 for heritability of 0.7, in a simulation study with popcorn. In this study, with real PE data, accuracy values of 0.75 for a heritability of 0.62 were obtained by strategy GEN + PhEN. It can therefore be inferred that the results are coincident, in view of the similarity in the observed proportions. According to (3), accuracy values 11 of ≥ 0.70 are desirable, i.e. the values of > 0.75 found for the studied traits indicate that satisfactory genetic gains can be obtained by GS.

### Genetic and environmental correlation and genetic gain

In this study, the correlations between the environments were high for most traits, mainly GY and PV, and EH and PH. According to (31), the environment becomes the cause of correlations when two traits are influenced by the same variations of environmental conditions, when negative values of this correlation show that the environment favored one trait at the expense of the other, and positive values, when both were benefited or harmed by the same causes of environmental variations.

The correlations between the trait pairs PH – EH and GY - PV were high and positive. Of course, the plants with highest EH tend to be the tallest plants (48). According to (49), these associations may allow indirect gains by selection in correlated traits.

The superiority of the strategy GEN over PhEN and GEN + PhEN agrees with the findings of (6,50,51), who suggested that the potential of GS is superior to that of phenotypic selection. However, this depends on a fitted prediction model, to be used in one or more RS cycles without recalibration. According to (52), the success of this strategy depends to a large extent on the maintenance of significant accuracy values, when the selection candidates are separated by one or more cycles of the training generation of the model.

According to (51), it is not possible to determine after how many cycles the model will become effective, because changes in the LD pattern, allele frequencies and polymorphism losses are unpredictable. According to (4), it is claimed that GS can be used from one to three RS cycles without a phenotypic evaluation, although this statement will need further evaluation. On the other hand, (41) performed five GS cycles and observed that the genetic gains in maize grain yield generally agreed with the predicted level, although the gains after the first cycle were unstable.

The results of strategy PhEN + GEN were slightly better than those of PhEN, due to the fact that strategy PhEN ignores the marker data to establish the kinship matrix, since for UENF-14 the kinship matrix is the proper identity (*G = I*). The population UENF-14 is unstructured, due to the recombination it underwent, with the progenies selected in the RS cycle, immediately before this study. According to (44) the closer the actual genetic relationship between the training and validation populations, the fewer markers will be required to achieve a satisfactory prediction accuracy.

The predicted annual gains for GY and PE by strategy GEN were much higher than those reported in previous RS studies for this population, where selection was performed exclusively with the evaluation of phenotypic family means (37,53–59).

Thus, it may be inferred that recurrent genomic selection results in high genetic gains, provided that: i) phenotyping is accurate, ii) the selection intensity is explored with the genotyping of several plants, increasing the number of selection candidates; and iii) GS is used for early selection in RS, thus reducing the interval between cycles, to spare the progeny test. However, more studies are needed to assess for how many subsequent RS cycles a genomic prediction model can be applied to determine GEBV with reasonable accuracy.

## References

1. Hallauer AR, Carena MJ, Miranda Filho JB. Quantitative Genetics in Maize Breeding. 3rd ed. Springer, editor. New York; 2010.

2. Borém A, Miranda GV, Fritsche-Neto R. Melhoramento de plantas. 7th ed. UFV, editor. Viçosa; 2017.

3. Resende MDV de, Resende Júnior MFR, Aguiar AM, Abad JIM, Sansaloni AAMC, Petroli C, et al. Computação da Seleção Genômica Ampla (GWS). EMBRAPA Florestas. Colombo; 2010.

4. Fritsche-neto R, Resende MDV De, Miranda GV, Dovale JC, Deon M, Resende V, et al. Seleção genômica ampla e novos métodos de melhoramento do milho. Rev Ceres. 2012;59(6):794–802.

5. Albrecht T, Auinger HJ, Wimmer V, Ogutu JO, Knaak C OM. Genome-based prediction of maize hybrid performance across genetic groups, testers, locations, and years. Theor Appl Genet. 2014;127(6):1375–86.

6. Yabe S, Hara T, Ueno M, Enoki H, Kimura T, Nishimura S, et al. Potential of Genomic Selection in Mass Selection Breeding of an Allogamous Crop: An Empirical Study to Increase Yield of Common Buckwheat. Front Plant Sci. 2018;9:1–12.

7. Xu S, Zhu D, Zhang Q. Predicting hybrid performance in rice using genomic best linear unbiased prediction. Proc Natl Acad Sci. 2014;111(34):12456–61.

8. Zhao Y, Mette MF, Reif JC. Genomic selection in hybrid breeding. Genomic Sel Crop Improv New Mol Breed Strateg Crop Improv. 2017;10:149–83.

9. Heffner EL, Lorenz AJ, Jannink JL, Sorrells ME. Plant breeding with Genomic selection: Gain per unit time and cost. Crop Sci. 2010;50(5):1681–90.

10. Daetwyler HD, Calus MPL, Pong-Wong R, Campos G de los, Hickey JM. Genomic prediction in animals and plants: Simulation of data, validation, reporting, and benchmarking. Genetics. 2013;193(2):347–65.

11. De Los Campos G, Hickey JM, Pong-Wong R, Daetwyler HD, Calus MPL. Whole-genome regression and prediction methods applied to plant and animal breeding. Genetics. 2013;193(2):327–45.

12. Valente MSF, Viana JMS, de Resende MDV, Silva FF, Lopes MTG. Genomic selection for plant breeding with different population structures. Pesqui Agropecu Bras. 2016;51(11):1857–67.

13. Neves LG, Davis JM, Barbazuk WB, Kirst M. A High-Density Gene Map of Loblolly Pine (*Pinus taeda* L.) Based on Exome Sequence Capture Genotyping. G3 Genes|Genomes|Genetics. 2014;4(1):29–37.

14. Purcell S, Chang C. Plink, Version 1.9. 2015.

15. Hill WG, Weir BS. Variances and covariances of squared linkage disequilibria in finite populations. Theor Popul Biol. 1988;33(1):54–78.

16. Endelman JB. Ridge Regression and Other Kernels for Genomic Selection with R Package rrBLUP. Plant Genome J. 2011;4(3):250–5.

17. Meuwissen THE, Luan T, Woolliams JA. The unified approach to the use of genomic and pedigree information in genomic evaluations revisited. J Anim Breed Genet. 2011;128(6):429–39.

18. Ertl J, Legarra A, Vitezica ZG, Varona L, Edel C, Emmerling R, et al. Genomic analysis of dominance effects on milk production and conformation traits in Fleckvieh cattle. Genet Sel Evol. 2014;46(1):1–10.

19. Misztal I, Legarra A. Invited review: Efficient computation strategies in genomic selection. Animal. 2017;11(5):731–6.

20. Misztal I. Inexpensive computation of the inverse of the genomic relationship matrix in populations with small effective population size. Genetics. 2016;202(2):401–9.

21. Butler D. asreml: asreml fits the linear mixed model. 2009.

22. R Core Team. R: A language and environment for statistical computing. R Foundation for Statistical Computing. Vienna; 2013.

23. Resende Jr MFR, Muñoz P, Resende MD V, Garrick DJ, Fernando RL, Davis JM, et al. Accuracy of genomic selection methods in a standard data set of loblolly pine (*Pinus taeda* L.). Genetics. 2012 Apr;190(4):1503–10.

24. Almeida Filho J de, Guimarães J, Silva F e, Resende M de, Muñoz P, Kirst M, et al. The contribution of dominance to phenotype prediction in a pine breeding and simulated population. Heredity. 2016;117(1):33–41.

25. Gilmour a R, Gogel BJ. ASReml User Guide. Release 20. 2007;

26. Meuwissen THE, Hayes BJ, Goddard ME. Prediction of total genetic value using genome-wide dense marker maps. Genetics. 2001;157(4):1819–29.

27. Erbe M, Pimentel E, Sharifi A, Simianer H. Assessment of cross-validation strategies for genomic prediction in cattle. In: 9th World Congress of Genetics Applied to Livestock Production: 2009. Giessen, Germany; 2010.

28. Bernardo R, Yu J. Prospects for genomewide selection for quantitative traits in maize. Crop Sci. 2007;47(3):1082–90.

29. Resende MDV De. Genômica quantitativa e seleção no melhoramento de plantas perenes e animais. Embrapa Floresta. Colombo; 2008.

30. Crossa J, Pérez-Rodríguez P, Cuevas J, Montesinos-López O, Jarquín D, Campos G de los, et al. Genomic selection in plant breeding : Methods, genomic selection in plant breeding: Methods, models, and perspectives. Trends Plant Sci. 2017;1–15.

31. Falconer D. Introdução a genética quantitativa. UFV. Viçosa; 1981.

32. Robertson A. Theory of limits in artificial selection. Proc R Soc London Ser B. 1960;234–49.

33. Terra T de F, Wiethölter P, Almeida CC de S, Silva SD dos A e, Bered F, Sereno MJC de M, et al. Genetic variability in maize and teosinte populations estimated by microsatellites markers. Ciência Rural. 2011;41(2):205–11.

34. Kashiani P, Ghizan S. Estimation of Genetic Correlations on Sweet Corn Inbred Lines Using SAS Mixed Model Pedram Kashiani and Ghizan Saleh Department of Crop Science, Faculty of Agriculture. Am J Agric Biol Sci. 2010;5(3):309–14.

35. Amaral AT do Junior, de Jesus Freitas IL, Guimarães AG, Maldonado C, Arriagada O, Mora F. Bayesian analysis of quantitative traits in popcorn (*Zea mays* L.) through four cycles of recurrent selection. Plant Prod Sci. 2016;19(4):574–8.

36. Pereira MG, Amaral Júnior AT. Estimation of Genetic Components in Popcorn Based on the Nested Design. Crop Breed Appl Biotechnol. 2001;1(1):3–10.

37. Santos FS, Amaral Júnior AT do, Júnior S de PF, Rangel RM, Pereira MG. Predição de ganhos genéticos por índices de seleção na população de milho pipoca UNB-2U sob seleção recorrente. Bragantia. 2007;66(3):389–96.

38. Rangel RM, Amaral AT, Scapim CA, Freitas SP, Pereira MG. Genetic parameters in parents and hybrids of circulant diallel in popcorn. Genet Mol Res. 2008;7(4):1020–30.

39. Schwantes IA, do Amaral Júnior AT, Gerhardt IFS, Vivas M, de Lima e Silva FH, Kamphorst SH. Diallel analysis of resistance to Fusarium ear rot in Brazilian popcorn genotypes. Trop Plant Pathol. 2017;42(2):70–5.

40. Heslot N, Jannink J-L, Sorrells ME. Perspectives for Genomic Selection Applications and Research in Plants. Crop Sci. 2015;55(1):1.

41. Combs E, Bernardo R. Accuracy of Genomewide Selection for Different Traits with Constant Population Size, Heritability, and Number of Markers. Plant Genome. 2013;6(1):2–7.

42. Lian L, Jacobson A, Zhong S, Bernardo R. Genomewide prediction accuracy within 969 maize biparental populations. Crop Sci. 2014;54(4):1514–22.

43. Zhang A, Wang H, Beyene Y, Semagn K, Liu Y, Cao S, et al. Effect of Trait Heritability, Training Population Size and Marker Density on Genomic Prediction Accuracy Estimation in 22 bi-parental Tropical Maize Populations. Front Plant Sci. 2017;8:1–12.

44. Liu H, Zhou H, Wu Y, Li X, Zhao J, Zuo T, et al. The Impact of Genetic Relationship and Linkage Disequilibrium on Genomic Selection. PLoS One. 2015;10(7):e0132379.

45. Muranty H, Troggio M, Sadok I Ben, Rifai M Al, Auwerkerken A, Banchi E, et al. Accuracy and responses of genomic selection on key traits in apple breeding. Hortic Res. 2015;2.

46. Lenz PRN, Beaulieu J, Mansfield SD, Clément S, Desponts M, Bousquet J. Factors affecting the accuracy of genomic selection for growth and wood quality traits in an advanced-breeding population of black spruce (*Picea mariana*). BMC Genomics. 2017;18(1):1–17.

47. Norman A, Taylor J, Tanaka E, Telfer P, Edwards J, Martinant JP, et al. Increased genomic prediction accuracy in wheat breeding using a large Australian panel. Theor Appl Genet. 2017;130(12):2543–55.

48. Cabral PDS, Júnior AT do A, Ismael Lourenço de Jesus Freitas R, Ribeiro M, Silva TR da C. Relação causa e efeito de caracteres quantitativos sobre a capacidade de expansão do grão em milho-pipoca. Rev Cienc Agron. 2016;47(1):108–17.

49. Cruz C, Regazzi A, Carneiro P. Modelos biométricos aplicados ao melhoramento genético. UFV. Viçosa; 2012.

50. Wong CK, Bernardo R. Genomewide selection in oil palm: Increasing selection gain per unit time and cost with small populations. Theor Appl Genet. 2008;116(6):815–24.

51. Jannink JL, Lorenz AJ, Iwata H. Genomic selection in plant breeding: From theory to practice. Briefings Funct Genomics Proteomics. 2010;9(2):166–77.

52. Müller D, Schopp P, Melchinger AE. Persistency of Prediction Accuracy and Genetic Gain in Synthetic Populations Under Recurrent Genomic Selection. G3 Genes|Genomes|Genetics. 2017;7(3):801–11.

53. Daros M, Amaral Júnior AT do A, Pereira MG. Genetic gain for grain yield and popping expansion in full-sib recurrent selection in popcorn. Crop Breed Appl Biotechnol. 2002;2(3):339–44.

54. Daros M, Junior AT do A, Pereira MG, Santos FS, Gabriel APC, Scapim CA, et al. Recurrent selection in inbred popcorn families. Sci Agric. 2004;61(6):609–14.

55. Freitas Júnior S de P, Amaral Júnior AT do, Rangel RM, Viana AP. Genetic gains in popcorn by full-sib recurrent selection. Crop Breed Appl Biotechnol. 2009;9:189–95.

56. Ribeiro RM, do Amaral Júnior AT, Gonçalves LSA, Candido LS, Silva TRC, Pena GF. Genetic progress in the UNB-2U population of popcorn under recurrent selection in Rio de Janeiro, Brazil. Genet Mol Res. 2012;11(2):1417–23.

57. Freitas ILJ, Júnior AT do A, Jr. SPF, Cabral PDS, Ribeiro RM, Gonçalves LSA. Genetic gains in the UENF-14 popcorn population with recurrent selection. Genet Mol Res. 2014;13(1):518–

58. Guimarães AG, Amaral Júnior AT do, Lima VJ de, Leite JT, Scapim CA, Vivas M. Genetic gains and selection advances of the UENF-14 popcorn population. Rev Caatinga. 2018;31(2):271–8.

59. Rangel RM, Teixeira A, Simões L, Gonçalves A, De S. Análise biométrica de ganhos por seleção em população de milho pipoca de quinto ciclo de seleção recorrente. 2011;473–81.

